# Maximal homology alignment: A new method based on two-dimensional homology

**DOI:** 10.1101/593228

**Authors:** Al Erives

**Affiliations:** Department of Biology, University of Iowa, Iowa City, IA, 52242-1324 USA

**Keywords:** *Drosophila*, enhancer DNAs, fractal repeats (FRs), gapped alignment (GA), microhomology, microparalogy, positional homology, replication slippage, tandem repeats (TRs), *ventral nervous system defective (vnd)*

## Abstract

Maximal homology alignment is a new biologically-relevant approach to DNA sequence alignment that maps the internal dispersed microhomology of individual sequences onto two dimensions. It departs from the current method of gapped alignment, which uses a simplified binary state model of nucleotide position. In gapped alignment nucleotide positions have either no relationship (1-to-None) or else orthological relationship (1-to-1) with nucleotides in other sequences. Maximal homology alignment, however, allows additional states such as 1-to-Many and Many-to-Many, thus modeling both orthological and paralogical relationships, which together comprise the main homology types. Maximal homology alignment collects dispersed microparalogy into the same alignment columns on multiple rows, and thereby generates a two-dimensional representation of a single sequence. Sequence alignment then proceeds as the alignment of two-dimensional topological objects. The operations of producing and aligning two-dimensional auto-alignments motivate a need for tests of two-dimensional homological integrity. Here, I work out and implement basic principles for computationally testing the two dimensions of positional homology, which are inherent to biological sequences due to replication slippage and related errors. I then show that maximal homology alignment is more informative than gapped alignment in modeling the evolution of genetic sequences. In general, MHA is more suited when small insertions and deletions predominantly originate as local microparalogy. These results show that both conserved and non-conserved genomic sequences are enriched with a signature of replication slippage relative to their random permutations.

---

Sequences of covalently-linked nucleotides and their evolutionary relationships are the basis of gene homology. The smallest possible scale to ascribe such homology is that of individual nucleotide positions (“site positional homology”). Of course this is possible only in the context of homological inference based on multiple nucleotide positions in a sequence.

By studying the function and evolution of regulatory DNA sequences (transcriptional enhancers in *Cionα* and *Drosophila*), which are not constrained by rigid, protein-coding, triplet reading frames (Erives *et al*. 1998; Erives and Levine 2000, 2004; Crocker *et al*. 2008; Erives 2009; Crocker *et al*. 2010; Crocker and Erives 2013; Brittain *et al*. 2014; Stroebele and Erives 2016), it has become apparent that site positional homology is mishandled by gapped alignment (GA). GA was originally adopted by necessity in order to compare protein sequences of divergent lengths (Braunitzer’s gappy comparisons of α- and β-hemoglobin chains), and later extended to nucleotide sequences (Bryson and Vogel 1965; Braunitzer 1966; Needleman and Wunsch 1970; Sankoff 1972). In retrospect, GA is not designed to model how regulatory nucleotide sequences evolve for the reasons to be explained here. This issue is exemplified by the fast-evolving regulatory DNAs of the speciose Hawaiian *Drosophila* (Brittain *et al*. 2014; O’Grady and DeSalle 2018). Nonetheless, this issue also is seen for the sequences encoding many important regulator proteins enriched in polyglutamine content and other repeats, including the Notch intracellular domain and sub-units of the Mediator complex (Tóth-Petróczy *et al*. 2008; Fuxreiter *et al*. 2008; Rice *et al*. 2015; Stroebele and Erives 2016; Erives 2017). As a related consequence, repeat alleles at these loci are not being accurately genotyped by modern genome assembly despite evidence of profound phenotypic effects (Rice *et al*. 2015; Chandler *et al*. 2017; Press and Queitsch 2017; Press *et al*. 2019).

Current techniques in computational molecular biology are agnostic as to the source of insertions and deletions (“indels”) and their homological relation to local sequence (Pevzner 2000). Consequently, GA does not treat an important source of genetic error that leads to alignment gaps: replication slippage (Strand et al. 1993; Haber and Louis 1998). Replication slippage is the result of DNA replication proceeding after a replication fork melts and re-anneals at an incorrect location thereby producing short duplicated stretches of sequence related through microhomology. Other enzymatic process, such as those that occur during repair or recombination, can generate similar errors (Haber 2000; Symington and Gautier 2011; Malkova and Ira 2013; Kowalczykowski 2015).

The type of microhomology that is ignored by GA is more accurately categorized as microparalogy because it is the result of local duplication. Microparalogy is relevant to the indel problem of GA, which attempts to identify where null characters should be inserted such that alignment is restored to homologous sequences. Thus the description of the goal of GA can be summarized as the restoration of uniform sequence lengths of one-dimensional strings by modeling sites as either orthologous or not, with orthology being used as a proxy for homology of all types. For want of an alternative approach to GA, it has not been possible to show how GA has impacted our understanding of sequence homology at the smallest scale. By explicitly addressing internal microparalogy, this study describes an alternative to GA that is biologically-grounded, computable, and productive in expected and unexpected ways.

One guiding proposition of MHA is that GA cannot find optimal alignments because it does not allow microparalogy to be captured into the same alignment columns. In short MHA changes previous assumptions of what criteria should be considered for optimality. Figure 1 gives a simple example as to how the long-standing search for a master equation for indel placement (Holmes 2017) may be side-stepped by shifting the overall approach and goals of DNA sequence alignment. In Figure 1 a trinucleotide repeat is variably expanded in two homologous sequences (Fig. 1A). Under GA these different sequences are aligned by the placement of null characters in any number of possible locations (Fig. 1B). This is not a contrived case because the majority of indels are the result of replication slippage, which is exacerbated within tandem repeats (TRs) (Ananda *et al*. 2013). Nonetheless, different homologs do not have to be of different lengths or even require the insertion of null characters *for GA to still be deficient*. The reason is as follows. Even in the gapped alignment of identical sequences that harbor various TRs, microparalogy remains de-localized over several columns despite their homological equivalence. This further highlights that GA is more focused on restoring length uniformity to character strings than modeling the evolutionary homology of genetic sequences.

**Figure 1.**
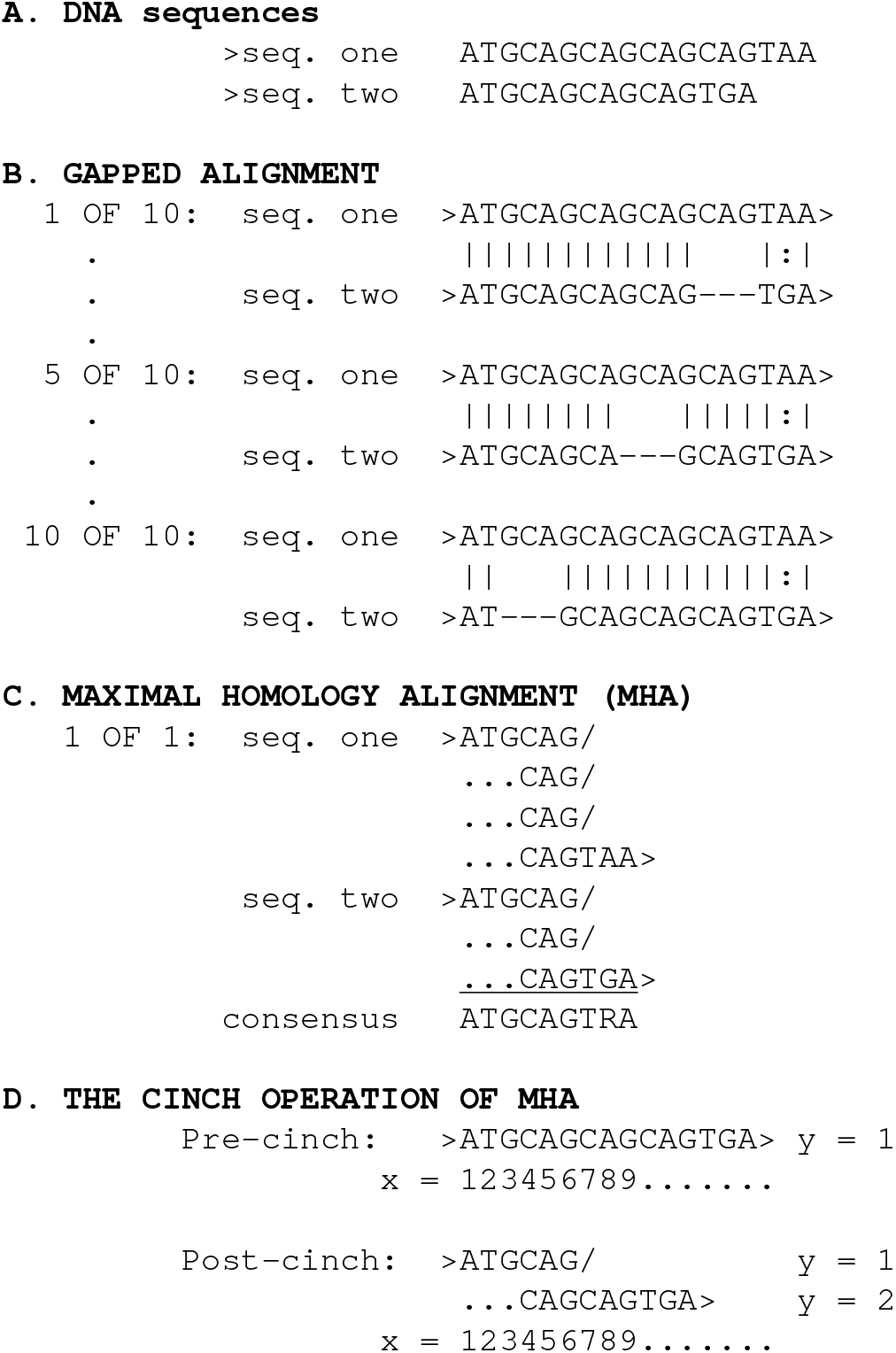
Gapped vs. maximal homology alignment. **(A)** The two homologous sequences shown will be aligned by GA (B) and MHA (C) for comparison. In the two homologs, a CAG trinucleotide repeat has variably expanded in number. **(B)** GA gives 10 possible alignments, differing by where a single parsimonious gap can be inserted in sequence two. There is no formal basis for favoring any one of these many alignments as optimal and any such choice would be artificial. All 10 gapped alignments are deficient in not encoding the multiple positions in microparalogical relationship with other nucleotide bases in the same sequence and in the other sequence. Thus GA artificially inflates the true number of columns of positional homology four-fold to 12 alignment columns. **(C)** Shown are the same sequences aligned by MHA, which are read left to right, top to bottom. In contrast to GA, MHA captures the micro-paralogy between CAG repeats into just one set of three columns. There is only one top-scoring alignment and there is no need for inserting null characters at all. Self-MHA treatments often produce 2-D objects that are automatically aligned without comparison to one another. The beginning and end of each sequence are marked with the greater-than symbol “>”, which is a convention to make it easier to see where different sequences begin and end in an MHA. The forward slash (“/”) is used as a symbol for an inferred replication slip, is sometimes referred to here as the slip symbol, and indicates the continuation of a (microparalogical) sequence on the next row. **(D)** MHA is based on the cinch operation. The cinch operation reduces the 2-D width along the *x*-axis while increasing its height along the *y*-axis. Dots are added to MHA sequences only to help visualize each alignment column, but otherwise carry no informational significance.

The presence of TRs is mutagenic with increasing repeat number and prone to elevating the local rates for substitutions and other errors (Amos 2010; Gemayel *et al*. 2010; Kelkar *et al*. 2011; Ananda *et al*. 2013; Duitama *et al*. 2014). Heterogeneous mechanisms likely underlie the instability at TRs, including the short microsatellite repeats (MSR, 1–9 bp units) and minisatellites (10–50 bp units) (Ellegren 2000a,b; Legendre *et al*. 2007). Within transcriptional enhancers some of this repeat content is built up of ancient repeats of repeats with dynamic instability (Crocker *et al*. 2010; Brittain *et al*. 2014). I refer to these more complex repeats as fractal-like repeats (FRs) because a fractal is a mathematical object which recursively features a repeating pattern at smaller scales. FRs have internal repeats of smaller unit sizes.

In summary there are two important considerations about evolving genetic sequences that this study seeks to address explicitly and precisely. The first consideration is that the internal microhomology of TRs and FRs is substantial enough to invalidate an implicit GA premise that indels are not typically homologically-relatable to local sequence. The second consideration is that nucleotide sequences composed of TRs and FRs are genetically unstable and self-amplifying, leading to a spectrum of repeat categories that include: (*i*) perfect, direct tandem repeats of a unit sequence (TRs); (*ii*) imperfect, direct tandem repeats harboring derived point substitutions but preserving the repeat unit length (“iTRs”); and (iii) fractal repeats where unit lengths are not preserved (both FRs and “iFRs”). The significance of the second consideration is that the MHA approach is a bigger challenge than simply identifying perfect non-overlapping TRs.

To address the shortcomings of gapped alignment in not considering local microparalogy, I began developing, formalizing, and implementing *maximal homology alignment* (MHA) to rescue microhomology that is unavoidably lost under GA [initial pre-print introducing MHA for the 2018 Drosophila Research Conference Erives (2018), and this study focusing on its principles]. Figure 1C shows how MHA rescues local microparalogy into the same alignment columns avoiding any need to insert null characters (dashes). The basic operation of MHA is the *cinch* operation, which is defined here as the lexical tokenization of a sequence into two paralogically overlapping sequences removed to separate rows (Fig. 1D). The cinch operation thereby cinches, or tightens, the two-dimensional width of a self-MHA, while increasing its height. Multiple sequence alignment (MSA) simply generalizes the cinch operation to entire disjointed homologs. The robust implementation of MHA presented here is able to (*i*) cinch identically perfect TRs, (*ii*) cinch imperfect TRs as determined by a substitution model and scoring system, (*iii*) cinch fractalized TRs in which repeats differ in unit size due to variation in number of sub-repeats, and (*iv*) resolve most overlapping but conflicting repeats of the various types.

In this study I focus on constructing and making practical use of a logical and systematic basis for MHA from biological considerations alone. In implementing a working version of MHA in a program called *maximal*, I discovered that necessary and sufficient computational tests for two-dimensional homological integrity can be derived from a few basic principles. The utility of these principles is that they show which desirable implications flow naturally without having to stipulate or test for them outright. One can mathematically and computationally test the dimensional integrity of homology with the tests described here. I then use MHA to characterize the extent of replication slippage in functional vs. non-functional biological sequences, and in *cis*-regulatory vs. protein-coding sequences. These explorations reveal the extent to which all biological sequences are imbued with a signature of persistent replication slippage and the extent to which there is negative selection to remove these errors from different logical compartments of the genome. In short the example of biological sequence can demonstrate how one dimension of position becomes two-dimensions informatically.

## Materials and Methods

### Statistics

Development of the *maximal* code base included a built-in Fisher-Yates shuffling option (Fisher and Yates 1953). Fisher-Yates shuffling was used to generate random permutations of natural sequences to use in millions of auto-MHAs over the course of software development to identify DNA strings that were problematic to cinch. Scripts to handle these randomization runs and to calculate width cinch ratios and related statistics from the data in the output log files are available with the code base.

The Fisher-Yates option was also used to run the randomization trials reported in Table 1 as controls. For these trials, the software was recompiled fresh each time to stipulate comparable sequence lengths suitable for comparisons to the test sequences (`FY_size`). Additional statistical tests are described in the footnotes to Table 1.

**Table 1.**
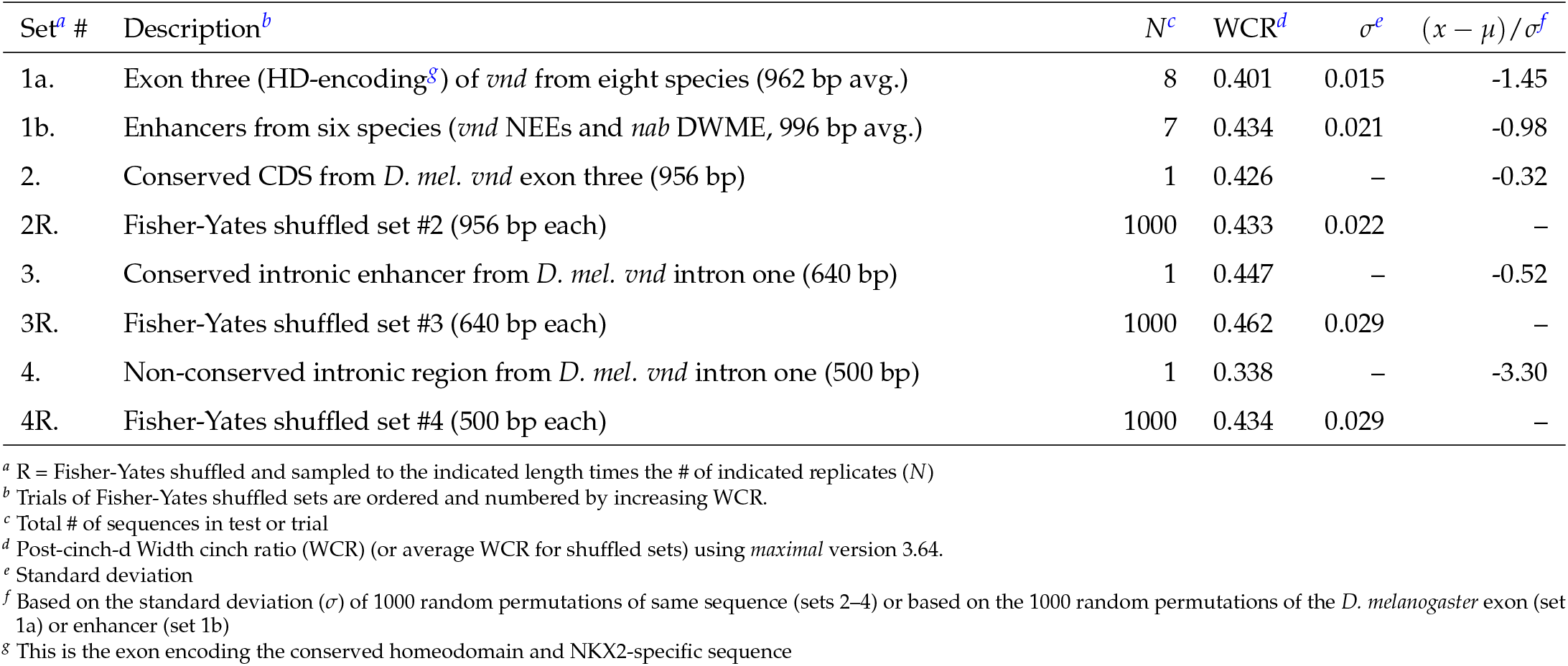
Width cinch ratios of biological vs. shuffled, and functional vs. non-functional sequences.

| Set<sup>a</sup> # | Description<sup>b</sup> | N<sup>c</sup> | WCR<sup>d</sup> | $\sigma^e$ | $(x - \mu) / \sigma^f$ |
| --- | --- | --- | --- | --- | --- |
| 1a. | Exon three (HD-encoding<sup>g</sup>) of *vnd* from eight species (962 bp avg.) | 8 | 0.401 | 0.015 | -1.45 |
| 1b. | Enhancers from six species (*vnd* NEEs and *nab* DWME, 996 bp avg.) | 7 | 0.434 | 0.021 | -0.98 |
| 2. | Conserved CDS from *D. mel. vnd* exon three (956 bp) | 1 | 0.426 | – | -0.32 |
| 2R. | Fisher-Yates shuffled set #2 (956 bp each) | 1000 | 0.433 | 0.022 | – |
| 3. | Conserved intronic enhancer from *D. mel. vnd* intron one (640 bp) | 1 | 0.447 | – | -0.52 |
| 3R. | Fisher-Yates shuffled set #3 (640 bp each) | 1000 | 0.462 | 0.029 | – |
| 4. | Non-conserved intronic region from *D. mel. vnd* intron one (500 bp) | 1 | 0.338 | – | -3.30 |
| 4R. | Fisher-Yates shuffled set #4 (500 bp each) | 1000 | 0.434 | 0.029 | – |

<sup>a</sup> R = Fisher-Yates shuffled and sampled to the indicated length times the # of indicated replicates (N)
<sup>b</sup> Trials of Fisher-Yates shuffled sets are ordered and numbered by increasing WCR.
<sup>c</sup> Total # of sequences in test or trial
<sup>d</sup> Post-cinch-d Width cinch ratio (WCR) (or average WCR for shuffled sets) using *maximal* version 3.64.
<sup>e</sup> Standard deviation
<sup>f</sup> Based on the standard deviation ( $\sigma$ ) of 1000 random permutations of same sequence (sets 2–4) or based on the 1000 random permutations of the *D. melanogaster* exon (set 1a) or enhancer (set 1b)
<sup>g</sup> This is the exon encoding the conserved homeodomain and NKX2-specific sequence

#### Algorithms

The following describes the initial two-dimensional array aspects of *maximal*, which is now supplemented with structure type definitions defined by the principles of internal twodimensional homology, which are described further below.

Path traversion: The *maximal* implementation of MHA begins similarly to the dynamic programming strategy used in global (Needleman and Wunsch 1970) and local (Smith and Waterman 1981) alignment with the construction of a path box, which when traversed describes the cinching required to produce local self-alignment. This path box is filled with scoring values according to a substitution matrix. Although the current *maximal* software recognizes the difference between DNA, RNA, protein, and alphabetic (non-biological) strings, a substitution model is only coded for DNA at the moment. I use a nucleotide substitution matrix that gives 1/2 the maximum identity score to transition substitutions. I also use a sparse matrix approach for efficiency and in particular find that we only need to fill in values in a hemi-diagonal band. A bandwidth of 200 bp is sufficient due to the known higher frequencies of tandem repeats (TRs) at increasingly lower *k*-mer sizes and their repeat number.

Cinch-t: In *maximal*, an initial computational module called “cinch-t” (t for *tandem repeats)* traverses a path in a manner completely unrelated to the trace-back strategies used by dynamic programming methods of traditional alignments. The cinch-t module uses a cinch-path finding strategy that begins in the upper left-hand corner and proceeds column by column from left to right, and from top to bottom in each column never passing the diagonal (hence the efficiency of a hemi-diagonal sparse matrix. This path will define the initial 2-D alignment. At column positions less than the bandwidth (the bandwidth defines the sparse matrix version of the path box), one begins at the first (top) row of each column, but after these initial columns one simply begins at the first intersection further below that is within the bandwidth. At each intersection with score *S* (*m, n*) > 0 for *n* > *m*, one evaluates whether the sum score of the diagonal beginning at that intersection and of length *k = n — m* surpasses a score threshold, which in our implementation is based on a *k*-mer dependent fraction of allowed transitions. Diagonals are currently discarded if they have a single non-transition mismatch, but different substitution strategies can be used. If a *k*-mer diagonal is found to surpass threshold, it is “cinched”, by which I mean that the line is terminated with an inferred replication slip character represented by the forward slash (“/”). Then the second or more repeats are placed under the first repeat unit block in a series of rows, one per additional TR unit, as microparalogical alignment. The slip character has no bearing on sequence length and is not meant to adjust spacing as the gap character does in GA. Instead the slip character merely conveys that the biological sequence continues on the next row below. Similarly, tick marks can be used to populate the upstream parts of rows that do not contain sequence merely as a visual guide for columnar alignment. These tick marks also do not have any bearing on alignment lengths. In the end, the cinch-t module cinches the first tandem repeats of smallest *k*-mer size even if they are constituents of a larger unit TR block of length *l* > *k* but then only cinches the larger *l*-mer TR blocks after that.

Other modules: In a subsequent module, called “cinch-k”, small TRs within the bigger *l*-mer repeats are then cinched like they were in the initial *l*-mer block. A cinch-l module tidies up long homopolymeric runs. Two additional modules (cyclelize and cinch-d) are described in the Results. An optional “relax-2D” module relaxes homopolymeric runs that did not contribute to successful cinch-d cinching operations. This module produces more proportional 2-D auto-alignments of single sequences, but cannot be optioned for pair-wise alignments so as to not make the cinch-d module non-productive.

#### Implementation

In the *maximal* implementation of MHA, TR unit sizes are limited by a hemi-diagonal sparse matrix approach with a bandwidth of 200 bp (in the *maximal* cinch-t module; see Materials and Methods). This is a reasonable cut-off because the majority of TRs and more extensive microsatellite repeats (MSRs) are composed of units that are less than 100 bp and most of this is less than 10 bp (Ananda *et al*. 2013). Thus in practice *x*-monotonicity is restored at a larger scale where a moving average of *x*-positions are taken for a window of a certain size.

Initial development versions of *maximal* used twodimensional arrays because these objects were easy to print computationally, and because a systematic principled basis for MHA was yet developed. There are several downsides to using two-dimensional arrays for MHA computation: they use a lot of memory, they do not store the one-dimensional positions for every two-dimensional point, and they are inefficient to crawl over while looking for sequence. More importantly, two-dimensional arrays do not take advantage of the principles of two dimensional homology that is described here.

The current C code has transitioned from exclusive use of two-dimensional arrays to also using structure type definitions, which can store one-dimensional positions alongside two-dimensional coordinates. The new structure type definitions allow for efficient testing of 2-D homological integrity (see The Principle of Continuity). A library of special functions (“check_tela”, “push_tela”, “get_1Dz”, “update_tela” and others) have been written and are in use (named for “tela” Spanish for a thin fabric, from Latin for thin web-like membrane). Future software versions will continue to expand the use of these data types and their special functionality for self-validation. For clarity, the structure `coord` array is called `stringy[i]` in *maximal’s* `main()` function but received and encoded as `tela[i]` in the library because the library functions can get calls from other structure arrays of the same `coord` type.

The github repository (https://github.com/microfoam/maximal.git) has a complete version of the code in the *maximal* implementation of MHA, as well as earlier versions of the code base. Below, pseudo-code is provided as a sketch of how the main principles of maximal homology alignment are implemented in structure type definitions of the C programming language. In this prototype, transition mutations are marked in the structure member `stringy[i].t` and derive from the equivalence symbols stored in the structure member `stringy[n].e`. Other linked structure members store *x* and *y* coordinates and base nucleotide characters. In this structure type definition, the structure array index *n* is the one-dimensional position, which as such requires no separate structure member to store. The “_tela” library functions are called with a simple pointer to the special structure `coord` type array as defined below.

~~~
DEFINE A STRUCTURE TYPE NAMED COORD:
typedef struct coord {
  int x IS COLUMN COORDINATE
  int y IS ROW COORDINATE
  char c IS CHARACTER SYMBOL: A, C, G, T
  char t IS FOR CALLED TRANSITIONS
  char e IS EQUIVALENCE CLASS: R,Y ETC. }
DECLARE A COORD-TYPE STRUCTURE ARRAY:
struct coord stringy[MALLOC SIZE]
CONFORM TO PRINCIPLE OF CONTINUITY:
If n = m+1, then one and only one case is true:
  stringy[n].x == stringy[m].x + 1 and
  stringy[n].y == stringy[m].y
or else
  stringy[n].y == stringy[m].y + 1 and
  stringy[n].x <= stringy[m].x
CONFORM TO PRINCIPLE OF EQUIVALENCE:
If stringy[n].x == stringy[m].x,
then stringy[n].t = stringy[m].t
~~~

### Data Availability

The *maximal* program, a working implementation of maximal homology alignment and application of the principles developed here, and its code base are all available on github (http://github.com/microfoam/maximal.git) under a GNU public license.

Also, available at the *maximal* github repository, is a large set of sequences identified as being initially refractory to cinching.

## Results

### A need for tests of 2-D homological integrity

Before defining the fundamental assumptions (principles) underlying two-dimensional positional homology and its implementation for MHA, I describe the types of computational problems that are solved in an MHA program called *maximal* (see Figure 2). Many of these problems were not obvious until implementation of MHA was tackled. The types of operations that one has to do often demand that the resulting 2-D objects be checked for two-dimensional homological “integrity”. A desire to check for this integrity then led to the identification and simplification of the principled logic of 2-D alignments.

**Figure 2.**
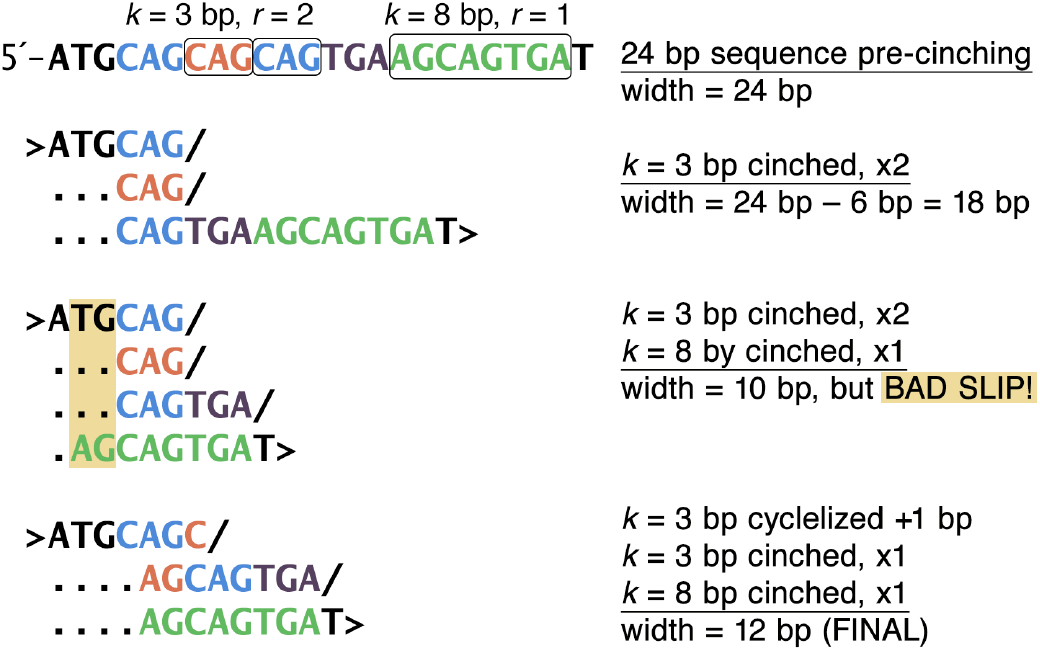
Overlapping repeats require special cinching for maintaining 2-D homological integrity. Shown is a 24 bp sequence containing two tandem repeat (TR) patterns that overlap. Both TRs are perfect in that the repeating units are exactly identical. (MHA in the *maximal* program also handles imperfect TRs.) The first repeat is a TR with a unit *k* = 3 size that repeats twice after the first unit (*r* = 2). When the two additional repeats are cinched under MHA, they end up being tucked underneath the first unit *k*-mer. Thus, 6 bp are tucked away to produce a 2-D auto-alignment that is 18 bp in width. If the second TR pattern of *k* = 8 is also cinched without consideration of the previously cinched upstream TR, then one would get a bad slip, which is characterized by alignment columns with non-homological calling (yellow shaded columns). For example, the T in the second position of the sequence erroneously gets called as being paralogous to the base nucleotide at position 16, which happens to be an A. Likewise, but less obvious, the G in position three gets erroneously called as being paralogous to the G in position 17. While these bases are identical, they should not be in the same column because there is no basis for inferring a paralogous relationship between them. The correct way of restoring local microparalogy into the same alignment columns is shown at the bottom. Here, the *k* = 3 repeat pattern is cyclelized as a repeat of CAG to a repeat of AGC in exchange for reducing the repeat number from *r* = 2 to *r* = 1 (see text for explanation of cyclelization operations). This allows the second *k* = 8 TR pattern to be cinched in a compatible manner for a final 2-D width of 12 bp. This final cinching represents a width cinch ratio (WCR) of 12/24 = 0.500. This also serves as an example of repeats of repeats, potentially created by one initial repeat’s instability.

#### Over slips and bad slips

Over slips are caused when a TR is cinched that overlaps a previously cinched TR. This results from a strategy of cinching sequence in a moving window that proceeds left to right (5′ to 3′). Over slips are particularly problematic when the earlier cinch effectively moves the upstream nucleotide bases of the second cinch to be downstream of the second cinches start along the *x*-axis. In the *maximal* program these are known as “bad slips” and are subsequently undone.

Bad slips are undone in the *maximal* program because the second cinch will always be larger than the first cinch due to the nature of the search. Over slips that do not need to be undone are simply ones that are compatible with the new cinch provided the new *x* starting coordinate is properly adjusted.

#### Cyclelization

*Cyclelization* is defined here as the operation of shifting the starting frame of a *cyclelizable* TR, which must necessarily have a minimum size of 2*k* + 1 where *k* is the TR’s unit size. A cyclelizable sequence would be something like 5’-`CAGCAGC`. In this sequence a repeating triplet can be cinched either as `CAG` at position four, or else as `AGC` at position five. In this example positions four and five are the starts of the *second* intact TR unit and where a new row would begin (*i.e*., a translational increment along the *y*-axis instead of along the *x*-axis). For this reason repeat number in this MHA implementation begins the count for the number of repeats only with a second unit repeat (*r* = 1, and the first unit could be considered repeat zero). This convention also avoids having to consider that every possible lexical tokenization of a sequence is a repeat of one unit.

In the case where there are no overlapping TRs, *maximal* simply takes the first frame. In the case of a bad slip, sometimes they can be cyclelized rather than undone, a preferred operation in regions of fractal repeats. In *maximal* this is handled by the “cyclelize” module, which currently cyclelizes *k*-mers of size two, which are the most common.

#### Fractal repeats (FRs)

As TR instability increases with repeat number, many areas of repeats show signatures of generating repeats of sub-strings of the founding repeats, which is referred to as fractal repeats here. The combination of the two overlapping repeats of Figure 2 is an example of the simplest type of FR or incipient FR. A more mature FR is shown in Figure 3. Nonetheless, sometimes there is a base pattern in which the unit size is different only because of a variable repeat number of an inner constituent repeat. Importantly, different unit repeat sizes are frequently normalized in size in the (*x*-axis) consensus of a 2-D self-alignment. Thus repeats of repeats can often be cinched by looking for tandem repeats in the consensus row and cinching accordingly if the operation would maintain 2-D integrity. In *maximal*, this is handled by the cinch-d module, which cinches *<u>de</u> novo* repeats of repeats.

**Figure 3.**
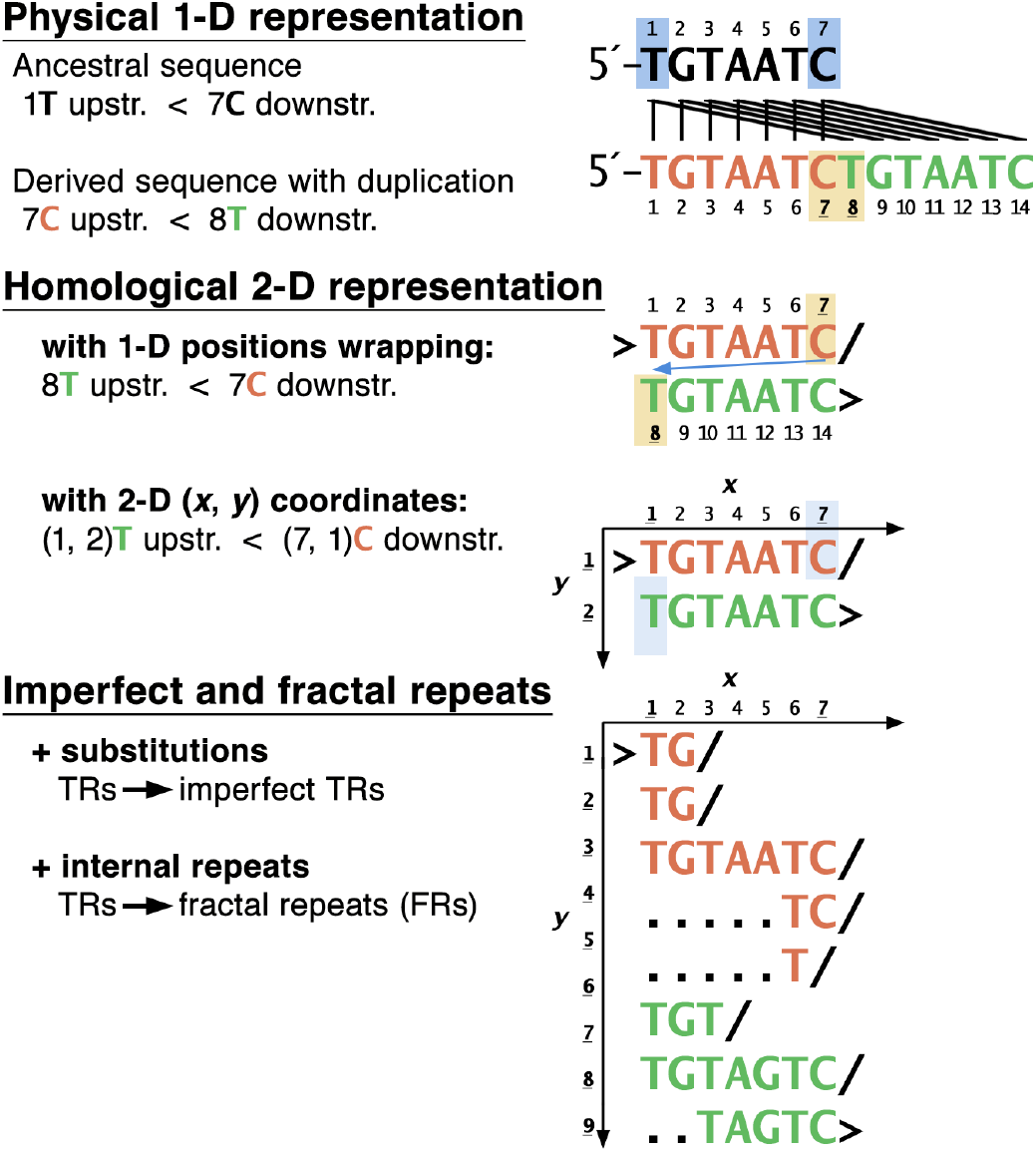
Dimensionality in biological sequences. Shown is an ancestral 7 bp sequence and one of its descendant homologs, which has inherited a duplicated tandem repeat of the ancestral sequence. The lines show the resulting 1-to-Many relationship between the ancestral sites and the descendant sites. These 1-to-Many relationships are not captured by gapped alignment. Note that in the physical representation of the 1-D sequence position seven occurs before position eight in the descendant sequence even though position eight is descended from position one in the ancestral sequence. In contrast, maximal homology alignment tucks one unit of the repeat underneath the columns of the first unit as shown. This MHA is shown once with 1-D numbered positions and again with 2-D coordinates for the *x* and *y* axes. Using the 2-D coordinates is more accurate and less confusing than using 1-D positions in an MHA. It is more accurate because each duplicated sequence inherits the same *x* coordinate positions spanned by the ancestral sequence. Thus this increased accuracy stems from the increased precision in modeling site position homology. The principles described in this study detail the logical relationships between 1-D positions and 2-D coordinates, and how they imply the sufficiency of a few computational tests to validate two-dimensional integrity of an MHA.

In short the possible operations of cinching both TRs and imperfect TRs have to deal with over slips, bad slips, cyclelizations, and FRs. Choosing which operations to do necessitates having available efficient tests of the validity of a proposed cinching operation. The following basic principles of MHA satisfy this need.

### The Principle of Continuity

The principle of continuity states how two adjacent nucleotide positions *m* and *n = m* + 1 from a one-dimensional biological sequence are related in the two dimensions that are required to model positional homology. I reserve the use of *m* and *n* to describe different one-dimensional positions, and reserve the use of *x* and *y* coordinates to describe the two-dimensional points of each one-dimensional position: *m* → (*x_m_, y_m_*) and *n* → (*x_n_, y_n_*). For clarity, I also reserve the use of the word “position” when referring to the unique one-dimensional location, and reserve the word “point” when referring to its (preferably unique) two-dimensional homological location, which is given in paired coordinates.

To describe the fundamental assumptions of MHA, I will also refer to biological homology mapping functions such as those that return either the *x* or *y* coordinates from the twodimensional representation given the one-dimensional position *n*. These mapping functions (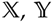, and 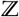) are discrete, nonlinear functions that map to and from the one-dimensional nucleotide space 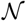 and the two-dimensional homological space 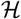 (see diagram of Equation 1 and Fig. 3). For clarity and ease of reference, additional bookkeeping functions 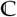 and 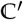 will refer to maps of the sequence indices represented by the spaces 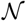 and 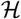 to the specific nucleotide bases in the sequence (A’s, C’s, G’s, and T’s). These points are summarized in Definition 0.1.

#### Definition 0.1.

The one-dimensional lattice space 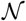 of indices of nucleotide position is related to the two-dimensional space 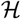 of homological indices via mapping functions 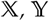, and 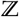. 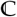 and 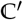 are the functions that map the spaces 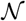 and 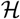, respectively, to the sequence B of nucleotide bases. These relationships are summarized by the diagram

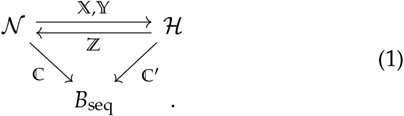

**Principle 1** (Continuity). Given one-dimensional molecular positions *m* and *n* = *m* +1, and 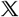 and 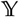 biological homology mapping functions, which return two-dimensional *x* and *y* coordinates respectively from one-dimensional positions, then one and only one of two cases can and must be true. Each of these cases is characterized by a pair of equations: Equations 2 + 3 (case one) or Equations 4 + 5 (case two). Case one corresponds to the non-homological relationship governed by:

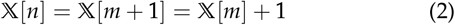

*and*

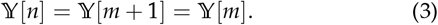

*Case two corresponds to the micro-paralogical relationship governed by:*

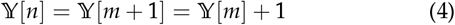

*and*

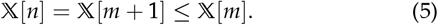

In short, adjacent nucleotide bases increase in homological position incrementally by single steps in either the *x* direction or the *y* direction, but never in both directions at once. Furthermore, when a position is incremented in the *y* direction 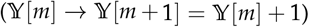, then the *x* coordinate must decrease or at least not increase 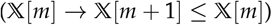. The second principle, the principle of paralogical equivalence, explains the condition under which this can occur.

The Principle of Continuity immediately implies four important features concerning homological space 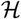: Implication 1.1: Mutual exclusivity of coordinate points, Implication 1.2: Existence of the ℤ mapping function, Implication 1.3: Guarantee of *y*-monotonicity, and Implication 1.4: Non-additivity of homological widths.

**Implication 1.1** (Mutual exclusivity of coordinate points). *Given one-dimensional molecular positions m and n such that n > m (or simply n ≠ m), then n and m can not share the same set of x and y homological coordinates*.

*Rationale*. For *n* > *m*, let *d* = *n* – *m*. For *n ≠ m*, let *n* be the larger of the two positions, and *m* be the smaller. From the principle of equivalence, it must follow that 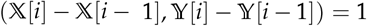. The reason for this is due to equations 2 and 4, which guarantee discrete one step movements in one axis from one nucleotide base to the next, and equations 3 and 5, which guarantee that when movement occurs in one axis, translational movement along the other axis is either zero or in the negative direction. Thus, if *d* = 0, then the *x* and *y* coordinates of *n* could not have changed from those of position m, and it must be that *n = m* (i.e., *n* and *m* cannot be different).

**Implication 1.2** (Existence of the ℤ mapping function). *By discretely counting the maximum of homological* 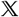 *and* 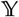 *increments relative to molecular increments, we can define a mapping function* ℤ *that corresponds to one-dimensional position. More succinctly, given a one-dimensional position, n, we can construct an equivalent homological mapping function based on two-dimensional coordinates*:

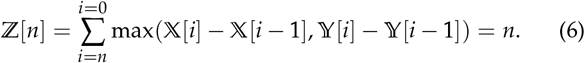

*Rationale*. From equations 2 and 3 under case one of nonhomology, we have that max(1,0) = 1. Alternatively from equations 4 and 5 under case two of microparalogy, we have that max(–*k*, 1) = 1, where — *k* < 0 and *k* is the *k*-mer size of the unit repeat, which is tucked underneath on the next row with a corresponding slip in the *x* axis. So by keeping a running count in this way we get a mapping function ℤ that behaves like a one-dimensional discrete position index but is based on two-dimensional biological homology (in conjunction with the Principle of Paralogical Equivalence).

With this definition of ℤ, we get a mapping function that counts two-dimensional homological position discretely, stepwise, and monotonically. This is something that we have taken for granted with nucleotide position, which always increases in discrete single steps and is monotonic in the 5′ to 3′ direction. This implication only holds if we respect the two mutually-exclusive cases described by the Principle of Continuity.

**Implication 1.3** (Guarantee of *y*-monotonicity). *Given onedimensional molecular coordinates m and n such that n > m, then*

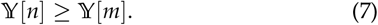

By omission this last implication implies that there can never be any guarantee of monotonicity for the *x*-axis. This result can be understood as follows. When reading DNA sequence left to right (5′ to 3′) one can eventually discover that a sequence eventually completes a second unit of a tandem repeat, resulting in a slipping of the *x*-axis position and a loss of monotonicity.

### MHA widths are not additive

The implications stemming from Principle 1 all apply to static MHAs, and so *y*-monotonicity and the absence of *x*-monotonicity hold only for static MHAs. In the *maximal* implementation of MHA, we have the case of cinch-t operations reversing bad slips when considering new sequence from a moving window for initial cinching. This is trivial example of an MHA addition operation, in which we are adding new un-cinched sequence to an existing (or growing) MHA. What follows is an important note about additivity of MHAs in general.

The absence of *x*-monotonicity indicates that twodimensional MHA width is not additive. The reasons for this have to do with 2-D width being a measurement of span along the *x*-axis, and with the nature of the absence of *x*-monotonicity. First, adding new sequence to a growing MHA might represent adding sequence that can be tucked under as additional microparalogy without having to change the existing rows. Second, adding new sequence might reveal a new completed repeat pattern at the edge, resulting in a smaller width for the existing MHA. Third, adding new sequence might reveal either a new set of cinching operation spanning the splice and/or a more optimal set of cinching operations that supersedes a previous set of operations. Thus, we can summarize the nature of MHA additivity with another implication:

**Implication 1.4** (Non-additivity of homological widths). Given two separate sequences a and b, and defining a + btobe the new longer sequence formed by linking a and b, then

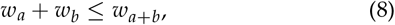

*where w_s_ is the two-dimensional (homological) width of sequence s of length n. In terms of the homological mapping function* 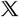, *the above relation is equivalent to* 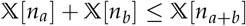. *Therefore, MHA widths are not additive*.

In the list of reasons leading to a new summed width that is less than the added widths, if *w_a_* is the width of the existing/growing MHA and *w_b_* is the width of the sequence being added to the 3′ end of *a*, then *w*_*b*1_ → *w*_*b*2_ < *w*_*b*1_ in the first and third examples, and *w*_*a*1_ → *w*_*a*2_ < *w*_*a*1_ in the second and third examples.

### The Principle of Paralogical Equivalence

By definition there are no orthological relations in an auto- or self-MHA, so internal homological relationships within a single sequence are all paralogical. The principle of equivalence states that two separate nucleotide bases can occupy the same alignment column (share the same *x* position) if and only if there is an assumption of paralogical equivalence (homology of position).

The principle of paralogical equivalence means that two onedimensional positions can share the same *x* coordinate if and only if the positions are paralogous positions in the two sequences located at 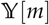 and 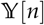. This means that first and foremost the two one-dimensional positions share identical *x*-coordinates. Second, it means that this is due to membership in a paralogous string, which must satisfy some simple probabilistic criteria that led to the calling of microparalogy in the first place. As such it is highly likely that the nucleotide bases are identical (*P*_1_) or else members of a preferred mutational equivalence class (*P*_2_ transitions: purine ⇔ purine and pyrimidine ⇔ pyrimidine). Nonetheless, the nucleotide bases may be different and belong to an improbable mutational class (*P*_3_ transversions: purine ⇔ pyrimidine), while being embedded in local microparalogy.

This principle states explicitly that we are establishing a primacy for the determination of the internal homology for a site position (an empty slot wherein one can place a nucleotide base) after considering the identity, similarity, or non-similarity of nucleotide bases in a column and nearby columns. Determination of site homology is of course dependent on sequence identity of neighboring bases.

**Principle 2** (Paralogical Equivalence). *Given one-dimensional molecular coordinates m and n > m, and the mapping function* 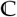, *which returns the nucleotide base from a one-dimensional position, if 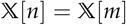, then*

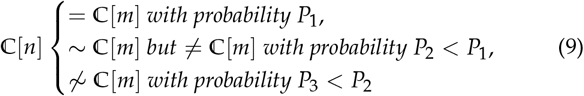

*such that P*_1_ + *P*_2_ + *P*_3_ = 1.

### A proposition for a “Principle of Precedence”

The principle of precedence is a proposition for relating and connecting maximal homology alignment to gapped alignment.

**Principle 3** (Precedence). *In aligning one-dimensional biological homologs, alignment proceeds first by auto-MHA (self-alignment), then by step-wise ordered multiple sequence alignment (MSA). During MSA additional cinching of divergent paralogy is first attempted. Relaxation of inferred micro-paralogy is attempted after that. Third and last the insertion of null characters (dashes) can be taken to finish the alignment if necessary. The insertion of null characters is reserved for columns containing unique non-paralogical sequence*.

*Rationale*. The purpose of maximal homology alignment is not to provide a master equation for indel placement in a gapped alignment. One could envision conducting gapped alignment and then considering local microparalogy at indels. However, the nature of microparalogy demands that all local sequence be considered, including sequences somewhat removed from the placement of a gap. This follows from the absence of a monotonicity guarantee in the *x*-axis unlike the situation for the *y*-axis (see Guarantee of *y*-monotonicity). Stated differently, the positions of gaps in a gapped alignment are not and can never be precise measurements of microparalogy.

It is expected that the cinching of inferred microparalogy might correspond to a false-positive cinching. Setting aside the possibility of determining the case of convergent evolution versus true homology, a false-positive cinch can occur when adjacent sequences independently evolve to be identical or nearly identical sequences. Under MHA, these sequences might be cinched as a type of false-positive microparalogy. Later, when conducting multiple sequence alignment, the position of false positive microparalogy might correspond to a place one would normally add null characters. However, because MHA is being employed, one has recourse to first relaxing the cinch and returning the cinched nucleotide bases to the would-be gap. Thus for pair-wise or MSA, over-cinching is inherently not an issue in MHA and can be reserved as a primary source of fill characters, relegating null characters as a second, backstop source. For single or auto-MHA, the *maximal* program is equipped with an optionable “relax-2D” function, which relaxes homopolymeric runs that did not aid in the cinching of fractalized TRs of different unit sizes. This is sometimes preferable for display purposes of MHAs of aesthetically-pleasing proportions.

### MHA mitigates the gap problem in divergent orthologs

Figure 4 shows the process of pairwise MHA for the edge of a conserved enhancer where the homology of sequence becomes difficult to discern in distantly-related *Drosophila* species. These are sequences for *D. melanogaster* and *D. erecta*, which began diverging from a common ancestor in the melanogaster subgroup about 10 million years ago (Powell 1997). The enhancer edge corresponds to the upstream flanking region of the NEE of the *ventral nervous system defective (vnd)* locus (see Fig. 5). For comparison standard global alignment-type GA and a genome browser reference alignment are shown to demonstrate the issues addressed by MHA (see Fig. 4A and 4B and legend). Self-MHAs of each sequence with normal non-aggressive MHA parameters are almost auto-aligned, while capturing microparalogy into the same columns (yellow highlighted sequences in Fig. 4C). Under the proposed principle of precedence, pair-wise alignment first attempts to restore alignment by cinching divergent microparalogy. Divergent microparalogy would be sequence that failed to meet a threshold alignment score during self-MHA, but which is evidently microparalogical when its homologous sequence from another species is able to surpass threshold in comparison. This first source of alignment (divergent microparalogy) is sufficient to restore perfect alignment, and avoids having to relax over-cinched false-positive microparalogy or insert any null characters (Fig. 4D). Figure 4D could be called the first pairwise DNA alignment under MHA.

**Figure 4.**
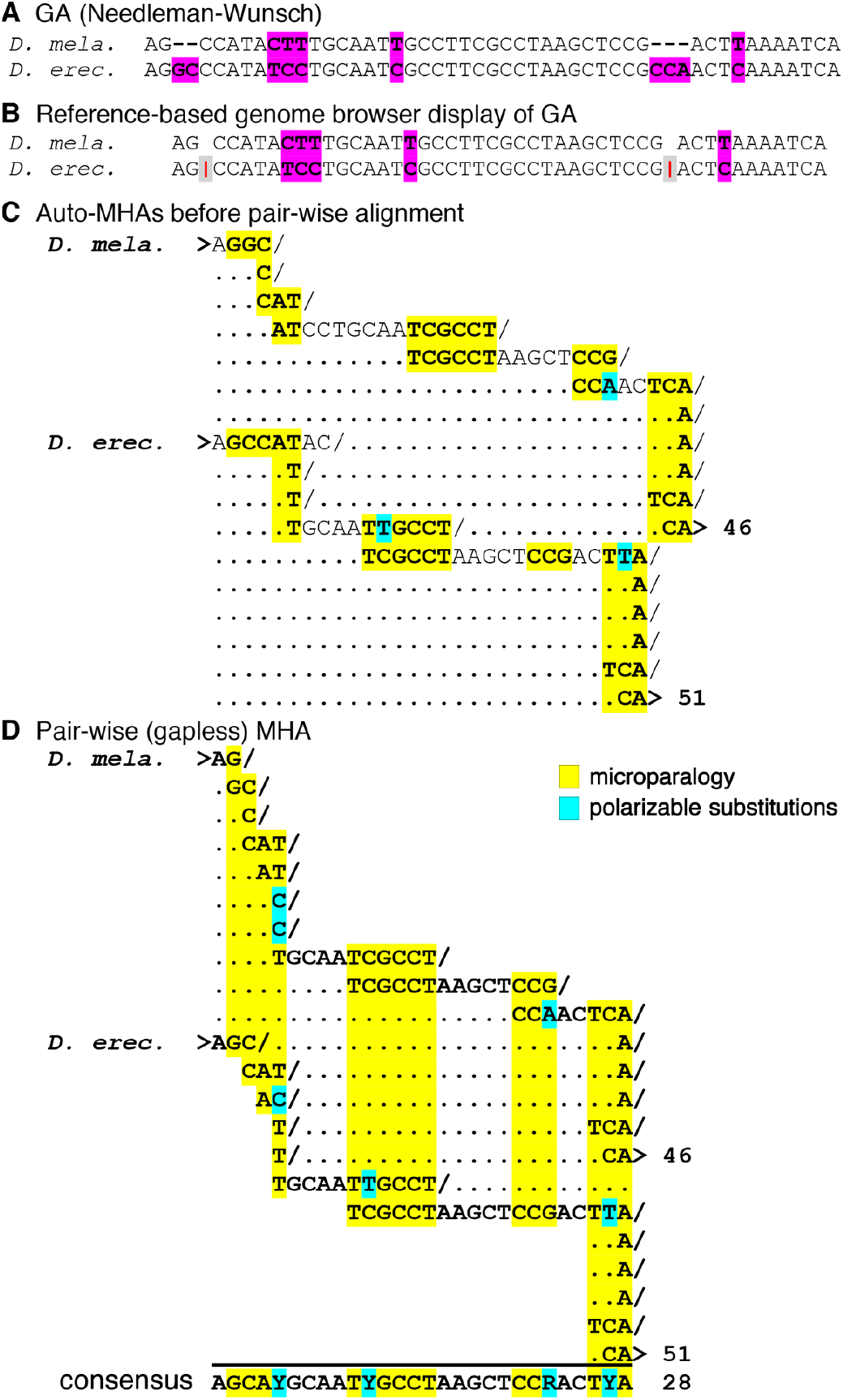
MHA does not typically need gaps to align distant homologs of different lengths. **(A)** Example GA using global alignment (Needleman and Wunsch 1970). Sequences correspond to the upstream edges of the *vnd* NEE in *Drosophila* species that began diverging 10 Mya. This region is where it becomes difficult to recognize homology. Differences (highlighted in magenta) are unpolarized in the absence of outgroup sequences, an issue that is often not manifested in MHA when it occurs in microparalogy. **(B)** Example of the same window of sequence in a genome browser view from the perspective of the reference species *D. melanogaster*. Sequences present only in *D. erecta* cannot be shown in relation to the reference genome, and is an issue that is not manifested in MHA. **(C)** Normal self-MHA preparation of the enhancer edges is almost sufficient to bring the divergent sequences into alignment. More aggressive self-MHA preparation can produce perfect auto-alignment but is sometimes problematic at FR “knots”. **(D)** Pair-wise alignment of the auto-MHAs. This alignment is the result of cinching divergent microparalogy in comparison to the homology (see text). The cyan highlighted sites are polarized substitutions based on the pair-wise alignment and later confirmed by consulting outgroups. These regions of microparalogy (yellow highlighting) are not identified in GA. This figure is modified from an earlier version that features slightly different alignment outcomes (Erives 2018).

Unlike GA this pairwise MHA highlights locations of microparalogy, which are more susceptible to instability and evolutionary divergence (see yellow highlighted columns in Fig. 4). These regions harbor all of the inferred substitutions, which are readily polarized *without* the aide of a third outgroup homolog (see cyan highlighted letters in Fig. 4). This is possible because the internal microparalogical dimension provides multiple (≥ 2) sequences for comparison. Altogether two-dimensional MHAs encode a much greater amount of inferred evolutionary genetic information than is possible in GA.

### 2-D widths of genetic sequences vs. random permutations

One can define a width cinch ratio (WCR) as the ratio of the final MHA cinch width (the 2-D width) to the starting sequence length (*i.e*., the 1-D width). *A priori* it is not clear how biological versus non-biological sequences would compare in their WCR values. On the one hand, biological sequences might be expected to have lower WCRs relative to randomly shuffled versions of the same sequences. This would be the case if the natural sequences are imbued with a deep signature of persistent replication slippage. On the other hand, it is possible that this signature is under intense negative selection. Tandem repeats and the fractal-like repeats of repeats are susceptible to instability and could be a target of persistent negative selection. If this latter case is likely, then we should be able to test this hypothesis by comparing functional biological sequences to non-functional biological sequences.

Table 1 shows the WCRs for a set of natural sequences (protein-coding exons or CDS’s, and developmental enhancers) from eight different species of *Drosophila*. These species are disparately related within the Sophophora subgenus of *Drosophila* and care was taken to remove closely-related sister species (sequences and scripts to run these tests are available with the *maximal* code repository.) In comparison to these sets of natural sequences (exons or enhancers), which have average WCRs of 0.401 and 0.434, respectively, randomly permutated sequences produced by Fisher-Yates shuffling of their *D. melanogaster* counterpart (exon or enhancer as appropriate) have much larger WCRs. The *Drosophila* exon set 1a is about –1.5 standard deviations below the average of 1000 permutations of the *D. melanogaster vnd* exon. Similarly, the *Drosophila* enhancer set 1b is about –1.0 standard deviations below the average of 1000 permutations of the *D. melanogaster vnd* enhancer.

To understand whether there is a basis for negative selection purifying functional sequences of replication slippage, I compared the upstream, non-conserved intronic region to the neurogenic ectodermal enhancer (NEE) of the *vnd* gene of *D. melanogaster*. Unlike the larger enhancer set 1b from several species, which encompasses the NEE and potentially extensive non-functional flanking sequences from different species with differently sized genomes, for this computational test I made the most contrasting comparison possible. I took the core 640 bp window of the NEE that is the most conserved across insects and compared it to the 500 bp window just upstream that is the least conserved (see Fig. 5). This 500 bp window is the largest obtainable relatively non-conserved stretch I could get without getting into the upstream exons or the downstream NEE. Table 1 shows that the non-functional intronic window has a WCR value (0.338) that is about 11% tighter than the core NEE (0.447), which occupies the same intron. In comparison, the protein-coding CDS of the third and terminal exon from the same locus, has a WCR that is smaller but more comparable to that of the functional (intronic) enhancer.

**Figure 5.**
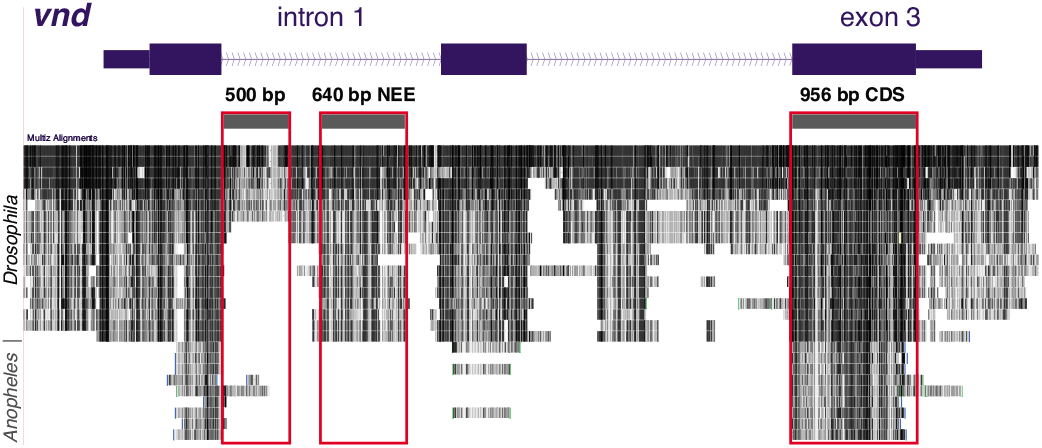
The *vnd* locus of *Drosophila melanogaster* with the three regions (red boxes) used in comparing functional vs. non-functional, and regulatory vs. protein-coding sequences. A 500 bp non-conserved intronic fragment was used as a proxy for a non-functional sequence relative to the conserved neurogenic ectodermal enhancer (NEE), which is located just downstream in the same intron. Specifically, a 640 bp core window of highly conserved sequence was used for the NEE. For comparison, a 956 bp protein-coding exon constituting the entire CDS of exon 3 was used. This region encodes the highly conserved DNA-binding homeodomain and NK2-specific domain of the Vnd transcription factor. Shown below are plots of the conservation across different *Drosophila* and *Anopheles* species based on the Multiz alignment blocks (*Blanchette et al*. 2004). Table 1 details how conservation in each of the three windows is correlated with larger (less pronounced) width cinch ratios (WCRs). Nonetheless, all three boxed regions are enriched in signatures of replication slippage relative to randomly shuffled sequences derived from each box.

I also performed multiple trials of Fisher-Yates permutations based on each of the three windows, separately (x1000 each). Each of these permutations is thus matched by sequence composition and length to their biological counterparts. Comparison to the non-biological shuffling experiments shows that, the biological windows tested from the *vnd* locus are 0.32 (CDS), 0.52 (NEE), and 3.30 standard deviations below the average of their matched trials. This suggests that while all of the windows are imbued with some degree of replication slippage signature relative to matched random permutations, the degree is correlated to the (relative absence of) evolutionary conservation and/or constraint for that window.

These results also underscore the effect on WCR ratios in the small *D. melanogaster* genome relative to other species in the genus (Gregory and Johnston 2008). For example, the *D. melanogaster vnd* exon 3 has a WCR of 0.433, while the Sophophora average is 0.401, representing a five-fold increase in the number of standard deviations below the average from random shuffling (see Table 1). Thus, MHA cinching occurs to a much greater extent in the orthologous sequences from other *Drosophila* species, which tend to have larger genomes.

## Discussion

The MHA Principle of Continuity and its implications together with the Principle of Paralogical Equivalence constitute the analogs to properties that we take for granted as being inherent to one-dimensional sequence, but which are not tenable in the two dimensions of biological sequence homology (see Fig. 3). This study thus challenges the assumption that an important goal in computational molecular genetics is to build 1-to-1 alignments of molecular sequence. Instead, I propose our goal should be to build alignments of positional homology, which must encompass both orthology and internal paralogy to be biologically relevant. The basic principles of maximal homology alignment are offered here as a foundation for biologically-relevant sequence alignment in way that is not possible in GA.

In the proposed Principle of Precedence, I argue for the use of “cinch-able” microparalogy and “relax-able pseudo-paralogy” as the primary alignment operations, while relegating the insertion of non-biological null characters (dashes) as a secondary, backstop operation. As shown in this study, MHA can reduce the need to use any fill characters (both pseudo-paralogy and null characters) as it rescues the replication slippage errors that are primarily responsible for gaps under GA (Fig. 4). This initial assessment and proposal will have to be tested with more extensive use of MHA and various implementations of MHA.

By working out the principles of two-dimensional sequence homology, I identify computational tests that are necessary and sufficient for validating two-dimensional homological representations and their underlying cinching operations (for example, see C pseudo-code in Materials and Methods, or functional C code in the repository for the *maximal* software project). For example, we would expect that different nucleotides in a DNA sequence should occupy different two-dimensional points in a 2-D auto-alignment. Initially, I thought this would be its own principle to be required and tested, a principle of exclusivity. But later, I saw that it was only an implication following from the principle of continuity along with the principle of *y*-monotonicity. So rather than have to test these three properties separately, one only has to test the main principle of continuity with the aid of the principle of paralogical equivalence.

In this first study, I used the principles to complete the first working version of *maximal* and to show that biological sequences are imbued with a signature of replication slippage that is under negative/purifying selection with increasing functional constraint (Fig. 5 and Table 1). Future versions of *maximal* will become much speedier because various operations in twodimensional arrays will be increasingly replaced in all places by the more abstract operations on the described C structure types that allow robust testing (see Materials and Methods). Currently this is done in the early cinch-t module, and its first application immediately reduced the *maximal* failure rate. The current version of *maximal* already uses both methods (two-dimensional arrays and structure arrays) for historical reasons and for software development tests. For example, special programming functions to check homological integrity and to translate 2-D coordinates to 1-D positions were written and applied.

### The evolutionary context of transversions

Rather than speculate on different possible ramifications of MHA, I simply state that its immediate direct use will be to model evolutionary divergence within a framework that respects the errors that characterize the enzymology of DNA replication. MHA is not designed nor suited for routine bioinformatics approaches such as retrieving similar sequences to a query sequence from a large database *(e.g.*, BLAST). To demonstrate the types of uses for which MHA might be deployed I pose an unusual but interesting MHA-motivated question concerning the context of transversion mutations.

Transitions are more common than transversions, which are errors that change a purine to a pyrimidine or *vice versa*. For this reason Kimura first introduced a model of DNA mutation to track different rates for transitions vs. transversions (Kimura 1980). The question is whether some transversions possibly originate as replication slippage. This would be the case if some transversions are actually templated by local sequence. Identical residues between sequence flanking the apparent transversion and the template sequence could easily mask its origin as a replication slippage error involving multiple base pairs rather than a single base substitution. Figure 6 shows one possible example from closely related homologous edges of the NEE from *D. simulans* and *D. mauritiana* as revealed by MHA using *maximal*. These species began diverging an estimated 240,000 years ago (Kliman *et al*. 2000; McDermott and Kliman 2008; *Garrigan et al*. 2012).

**Figure 6.**
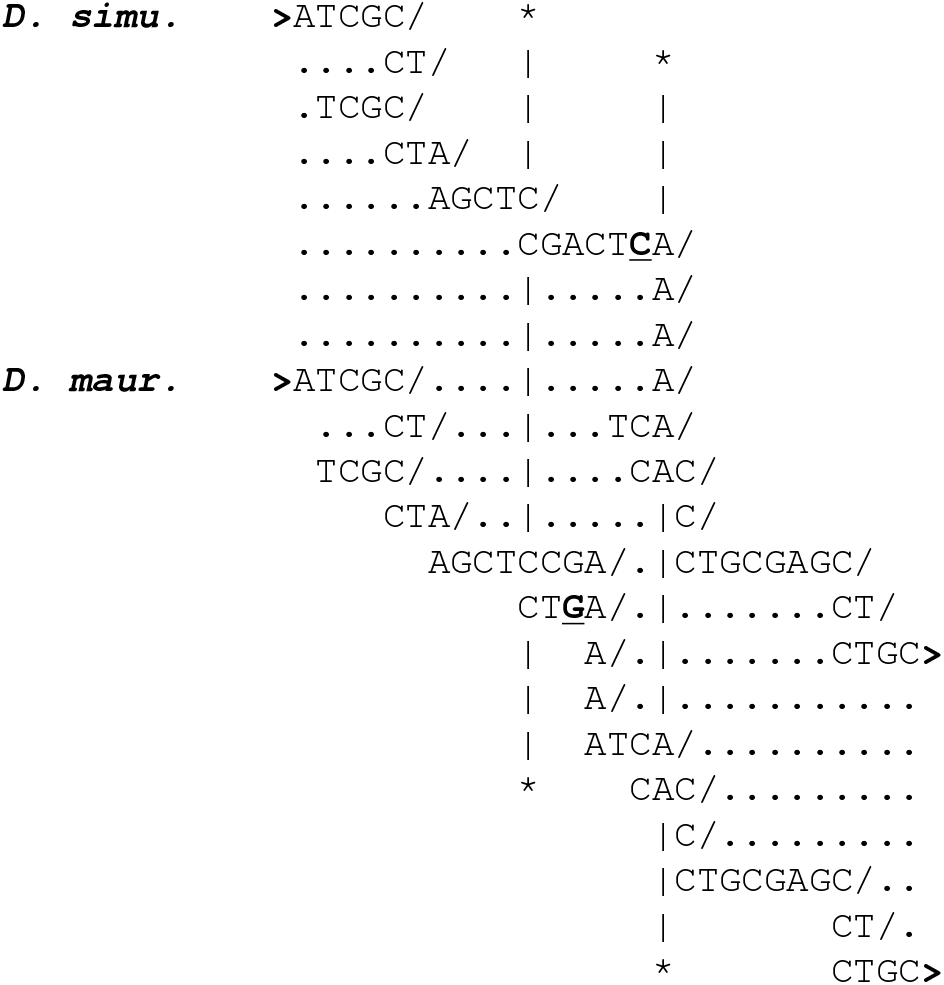
A transversion substitution potentially mistem-plated by canonical complementarity with local microparalogy. Shown are the raw maximal homology alignments produced by the *maximal* software. These sequences correspond to the upstream edges of the beginnings of the neurogenic ectodermal enhancers (NEEs) from *Drosophila simulans* (top sequence) and *Drosophlia mauritiana* (bottom sequence), which correspond to two closely related species that diverged < 250 kya. These sequences are also nested for compaction (a built-in feature of *maximal*). The MHA segment spanning the asterisked lines, inclusive of the lines, is the only segment where the cinching is different between the two sequences. This is the result of a single transversion mutation in *D. mauritiana* relative to other closely-related species: *C → G* (bold underlined letters). This transversion triggers the cinching of an imperfect tetranucleotide sequence with one transition mismatch. In the context of the MHA representation, it could indicate that the transversion substitution may correspond to a transition substitution that was mis-templated from adjacent microparalogy.

The two sequences in Figure 6 are identical except for a single transversion substitution. In comparison to outgroups (not shown), *D. mauritiana* is found to have a derived C → G transversion substitution (bold underlined G letter). Due to this substitution the auto-MHA process for the derived *D. mauritiana* sequence cinches up differently in the middle third of the alignment. This is due to the CTGA sequence forming an imperfect repeat with the upstream tetra-nucleotide CCGA. This might suggest that the apparent transversion may have been mistemplated and if so would represent microparalogy produced via complementary base-pairing. Thus, MHA may allow for the in-depth study of the sequence context surrounding transversion mutations and the extent to which transversions are correlated to adjacent microparalogical content. It will then be informative to see whether this is more common during DNA replication, meiotic recombination, and/or various repair pathways.

## Acknowledgments

This work was supported in part by an NSF CAREER award to AE to study morphogen gradient enhancer readouts (IOS:1239673), and an anonymous benefactor who provided support in part “to complete work on publishing at least one paper on molecular evolution”. I personally thank John Reinitz for his time in providing detailed comments on an earlier version of this manuscript and for suggesting the inclusion of a diagram (Definition 0.1) depicting the relationships between functions and spaces.

